# Profiling *APOL1* Nephropathy Risk Variants in Genome-Edited Kidney Organoids with Single-Cell Transcriptomics

**DOI:** 10.1101/780684

**Authors:** Esther Liu, Behram Radmanesh, Byungha H. Chung, Michael D. Donnan, Dan Yi, Amal Dadi, Kelly D. Smith, Jonathan Himmelfarb, Mingyao Li, Benjamin S. Freedman, Jennie Lin

## Abstract

**Background:** DNA variants in *APOL1* associate with kidney disease, but the pathophysiological mechanisms remain incompletely understood. Model organisms lack the *APOL1* gene, limiting the degree to which disease states can be recapitulated. Here we present single-cell RNA sequencing (scRNA-seq) of genome-edited human kidney organoids as a platform for profiling effects of *APOL1* risk variants in diverse nephron cell types.

**Methods:** We performed footprint-free CRISPR-Cas9 genome editing of human induced pluripotent stem cells (iPSCs) to knock in *APOL1* high-risk G1 variants at the native genomic locus. iPSCs were differentiated into kidney organoids, treated with vehicle, IFN-γ, or the combination of IFN-γ and tunicamycin, and analyzed with scRNA-seq to profile cell-specific changes in differential gene expression patterns, compared to isogenic G0 controls.

**Results:** Both G0 and G1 iPSCs differentiated into kidney organoids containing nephron-like structures with glomerular epithelial cells, proximal tubules, distal tubules, and endothelial cells. Organoids expressed detectable *APOL1* only after exposure to IFN-γ. scRNA-seq revealed cell type-specific differences in G1 organoid response to *APOL1* induction. Additional stress of tunicamycin exposure led to increased glomerular epithelial cell dedifferentiation in G1 organoids.

**Conclusions:** Single-cell transcriptomic profiling of human genome-edited kidney organoids expressing *APOL1* risk variants provides a novel platform for studying the pathophysiology of APOL1-mediated kidney disease.

**SIGNIFICANCE STATEMENT:** Gaps persist in our mechanistic understanding of APOL1-mediated kidney disease. The authors apply genome-edited human kidney organoids, combined with single-cell transcriptomics, to profile *APOL1* risk variants at the native genomic locus in different cell types. This approach captures interferon-mediated induction of *APOL1* gene expression and reveals cellular dedifferentiation after a secondary insult of endoplasmic reticulum stress. This system provides a human cellular platform to interrogate complex mechanisms and human-specific regulators underlying APOL1-mediated kidney disease.

## INTRODUCTION

APOL1-mediated kidney disease accounts for a portion of the excess risk of chronic kidney disease (CKD) and end-stage kidney disease (ESKD) among African American patients^1,2^. The *APOL1* high-risk genotype, defined as the presence of two risk alleles (G1 or G2 coding variants), increases the risk of developing CKD, but not all individuals with the high-risk genotype develop disease^3,4^. Much remains unknown regarding mechanisms and modifiers that render the disease incompletely penetrant, and complex interactions underlying these mechanisms are difficult to model outside *APOL1*’s native genomic locus. As such, current gaps in knowledge may not be fully addressed by induction of transgenic *APOL1* expression *in vivo* or *in vitro*. Additionally, because *APOL1* is widely expressed across different cell types, studying *APOL1* risk variants solely within a specific type of cell (e.g., podocytes) may not fully capture how these variants affect the kidney.

Human kidney organoids derived from induced pluripotent stem cells (iPSCs) can be utilized to model genetic disease mechanisms in the native genomic context and cell-type heterogeneity within the kidney^5–9^. Using CRISPR-Cas9 mediated genome editing, we engineered iPSCs homozygous for the G1 risk allele and differentiated these cells into three-dimensional kidney organoids. To evaluate cell-type specific effects of the *APOL1* high-risk genotype, we also performed single-cell RNA-sequencing (scRNA-seq), which we and others have previously leveraged to uncover novel biology of how cell-specific phenotypes contribute to kidney development or disease in organoids and other models^10–14^. Here we present the application of genome-edited iPSC-derived kidney organoids and single-cell transcriptomics to profile APOL1-mediated effects on kidney organoids relevant to disease processes.

## METHODS

### Induced Pluripotent Stem Cell (iPSC) Culture

iPSC lines previously derived from fibroblasts (Harvard Stem Cell Institute, 1016SevA) and peripheral blood mononuclear cells (WiCell, Penn134-61-26) were maintained in feeder-free culture on 10 cm dishes coated with 0.5% Geltrex (Gibco) in Modified Tenneille’s Special Recipe 1 (mTeSR1, STEMCELL Technologies), supplemented with 1% Penicillin/Streptomycin (Gibco) and 0.02% Plasmocin (Invivogen). iPSCs were confirmed to be mycoplasma-free and below passage 48. They were passaged using 1:3 Accutase (STEMCELL Technologies).

### CRISPR Cas-9 Genome Editing

*APOL1* G1 risk variants (rs73885319 and rs60910145) were introduced into the 1016SevA iPSC line through a genomic footprint-free approach (Figure 1A, Figure S1A)^15,16^. Briefly, the homology-directed repair (HDR) template containing the G1 variants was engineered using the MV-PGK-Puro-TK vector (Transposagen Bio), referred to as PMV vector, which houses a removable puromycin selection cassette flanked by two homology arms. The puromycin cassette is excisable by a piggyBac transposase, leaving only a “TTAA” sequence behind that can be seamlessly introduced into a coding sequence by carefully choosing sites where the change would be synonymous. The G1 variants were engineered by two-step PCR of G0 genomic DNA (gDNA) (Figure S1A, Table S1) to create the donor template for homology arm A, designed to flank the upstream portion of the puromycin selection cassette. Arm B, designed to flank the downstream end of the selection cassette, was amplified from G0 gDNA by traditional PCR. Both Arm A and Arm B underwent separate TOPO TA cloning reactions (Invitrogen) to insert into a stable vector for subsequent subcloning into the PMV vector. Stepwise sequential double restriction-enzyme digests and homology arm ligations were performed on the PMV vector with the following pairs of restriction enzymes: Not1-HF and Bbs1-HF, Nco1-HF, and Bsa1-HF (New England Biolabs). The ends of both homology arms bordering the cassette harbor the “TTAA” piggyBac transposase cut sequence, thus allowing for the transposase to excise the cassette from both ends and leave behind the “TTAA” sequence in a scarless fashion (Figure S1B). To make this genome editing event footprint-free, we selected a codon site that would allow the “TTAA” nucleotide sequence to be knocked in without altering the APOL1 amino acid sequence. We identified a leucine (an amino acid encoded by six different codons including “TTA”) flanked by an adenine to be the site of cassette entry and excision (Figure S1C). A guide RNA sequence with a suitable protospacer adjacent motif (PAM) was found nearby the excision site (Figure 1A, Table S1) and cloned into gRNA_Cloning Vector (Addgene 41824)^17^. The donor template incorporates a point mutation at the PAM site to destroy it after HDR to prevent recutting. iPSCs were then electroporated with the guide vector, hCas9 (Addgene 41815)^17^, and the G1-PMV donor plasmid (control lines were electroporated with the guide vector only). 48 hours later, 10 µg/mL puromycin was added to iPSC culture to select for cells expressing the donor plasmid (Figure S1A). Seven days later, iPSC colonies were evaluated for insertion of HDR template by PCR and Sanger sequencing validation. After expansion of the successfully knocked-in colonies into separate lines, the piggyBac transposase expression vector (Transposagen Bio) was introduced by electroporation. Additional screening of genotype was performed to validate the puromycin cassette excision.

**Figure 1.**
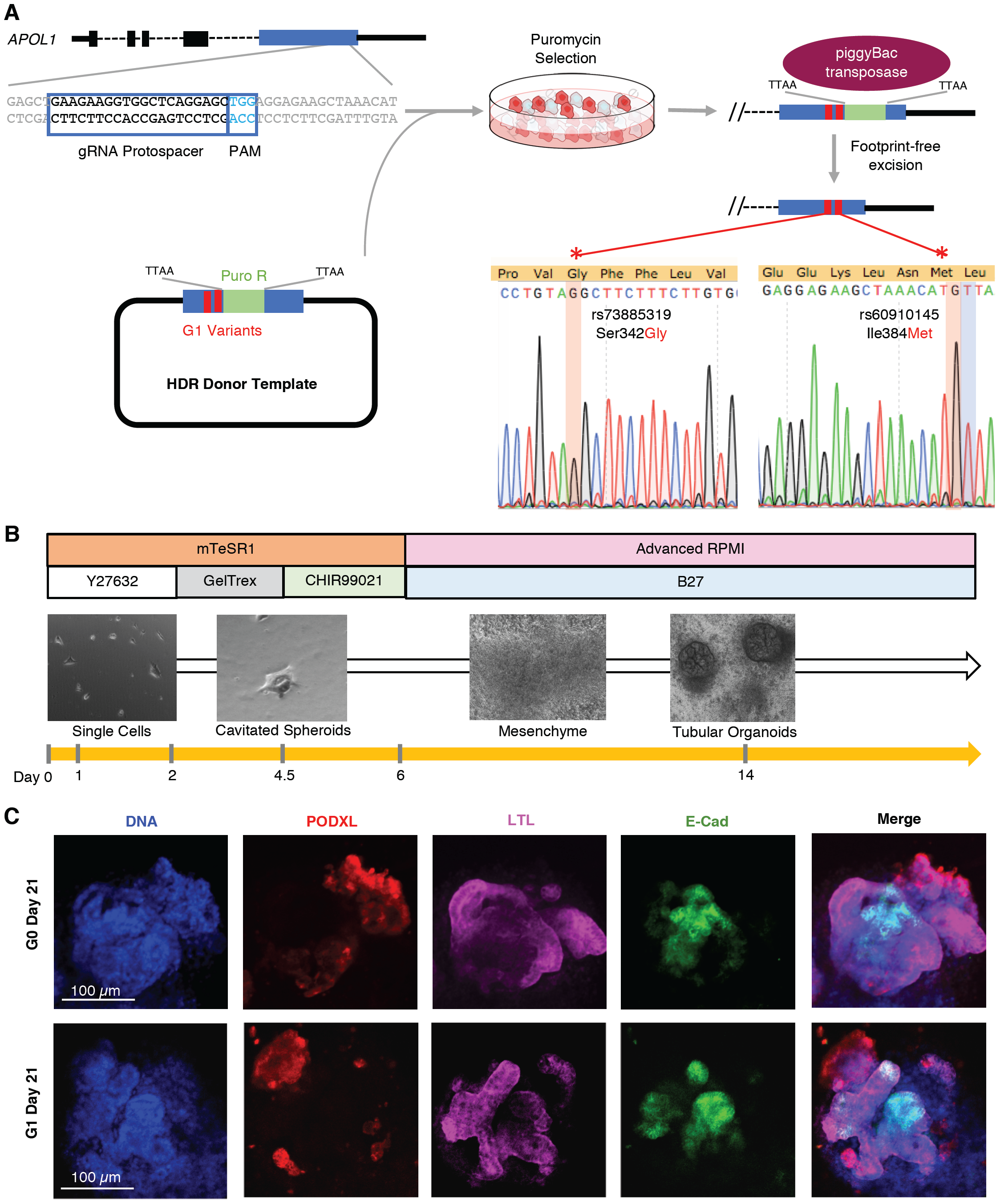
Engineering APOL1-G1 kidney organoids from iPSCs. **A.** Schematic summarizing the CRISPR-Cas9 approach to knocking in the G1 risk variants rs73885319 and rs60910145 into the 1016SevA line, including the chosen protospacer, HDR donor template design leveraging the piggyBac transposon system, and Sanger sequencing validation of successful variant knock-in and selection cassette excision. **B.** Overview of the Freedman kidney organoid differentiation protocol, with light microscopy of the 1016SevA line forming tubular organoids. **C.** Confocal immunofluorescence images of nephron markers in representative G0 and G1 organoids on day 21 of differentiation.

### iPSC-Derived Kidney Organoid Differentiation

iPSCs were differentiated into kidney organoids following the previously published Freedman et al. protocol^5^ (Figure 1B). Briefly, iPSCs were dissociated with 1:3 Accutase and plated onto 24-well plates pre-coated with 0.5% GelTrex in mTeSR1 supplemented with 10uM Y-27632 ROCK Inhibitor (STEMCELL Technologies). 24 hours later another layer of GelTrex at 1.5% was added in mTeSR1 media. At the end of the fourth day the medium was replaced with Advanced RPMI (Gibco) supplemented with 12 µM CHIR-99021 (STEMCELL Technologies). Approximately 60 hours later, the medium was changed to Advanced RPMI with B27 (Gibco). Organoids were cultured in this medium until collection at day 25.

### Induction of *APOL1* Expression and Endoplasmic Reticulum (ER) Stress

Day 24 G0 and G1 kidney organoids in identically plated wells of a 24-well plate were treated with interferon-gamma (IFN-γ, 25ng/mL, PeproTech) for 24 hours to induce *APOL1* expression. ER stress was induced by adding 5 µM Tunicamycin (Tocris) for 24 hours.

### Single-cell RNA Sequencing (scRNA-seq)

We performed scRNA-seq on G0 and G1 day 25 kidney organoids treated with vehicle, IFN-γ, or both IFN-γ and tunicamycin for 24 hours. Organoids were dissociated from the well with TrypLE Express (Gibco) and processed into single-cell suspension by gentle intermittent pipetting while incubating in a ThermoMixer (Eppendorf) for up to 15 minutes. Single-cell libraries were prepared using the 10x Genomics Chromium droplet-based platform and the Single Cell 5’ Library Construction Kit (10x Genomics), which was chosen to increase read coverage over the 3’ chemistry. At least three technical replicates were included in each prepared library, with targeted cell recovery of 4,000 cells per library. Libraries generated from three separate experiments were assessed for quality control, pooled, and sequenced on the NovaSeq 6000 (Illumina) through the University of Illinois Genomics Core. The libraries were processed for 150-base pair (bp) paired-end reads, at an average sequencing depth of 114,000 reads per cell.

### Immunofluorescence

Organoids were fixed in 4% paraformaldehyde for 15 minutes at room temperature. After fixing, samples were washed in PBS, blocked in 5% donkey serum (Millipore)/0.3% Triton-X-100/PBS for one hour at room temperature, incubated overnight in 3% bovine serum albumin (Millipore)/PBS with primary antibodies, washed, incubated with Alexa-Fluor secondary antibodies (Invitrogen) and DAPI, and washed into PBS for storage. Primary antibodies included PODXL (AF1658; R&D; 1:500) and ECAD (ab11512; Abcam; 1:500). Stains included fluorescein-labeled LTL (FL-1321; Vector Labs; 1:500). Fluorescence images were captured using an inverted Nikon epifluorescence Eclipse Ti or A1R confocal microscope.

### Real-Time Quantitative Polymerase Chain Reaction (RT q-PCR)

RNA was isolated from day 25 kidney organoids using the PureLink Kit (Invitrogen). RNA was then reverse transcribed using High-Capacity cDNA Reverse Transcription Kit (Applied Biosystems). RT-qPCR reactions for *APOL1* (Thermo Fisher, Hs01066280_m1) and *ACTB* (Hs01060665_g1) were run in duplicate on the QuantStudio 5 Real-Time PCR System (Applied Biosystems). Relative *APOL1* expression was calculated using the 2^-ΔΔCT^ method. Statistical significance was tested using two-way ANOVA in Graphpad (Prism).

### Immunoblot

Organoids were washed with PBS and lysed in RIPA buffer (Thermo Scientific) with protease inhibitor (Thermo Scientific) and benzonase nuclease (Thermo Scientific). Cell lysates were cleared by centrifugation for 10 minutes at 14,000 × g at 4º C. Proteins (20µg) were heated at 95°C with β-mercaptoethanol and then separated by SDS PAGE (Invitrogen). Nitrocellulose membranes were blocked in 5% nonfat milk in PBST for 1 hr and then incubated in primary antibody overnight at 4ºC, followed by incubation with HRP-conjugated secondary anti-Ig (Abcam; 1:5000). The primary antibodies used in this study were anti-APOL1 (HPA018885; Sigma; 1:1000), anti-β-actin (#4970; Cell Signaling Technologies; 1:1000). Immunoblot signals were developed under chemiluminescence (Thermo Fisher) and digitally imaged (BioRad).

### Bioinformatic Analyses

Approach to bioinformatic analyses of scRNA-seq data are presented in Supplemental Material.

### Data Availability

All scRNA-seq data have been deposited in Gene Expression Omnibus under accession number GSE135663.

## RESULTS AND DISCUSSION

Using CRISPR Cas-9 genome editing, we engineered an iPSC line homozygous for the *APOL1* G1 risk alleles, with G0 controls on an isogenic background from the same parent line. Unlike previous cell lines used to study risk-variant APOL1 through transgenic expression ^18–21^, this engineered line houses the G1 variant at its native genomic locus under the control of *APOL1’*s endogenous regulatory elements. To increase HDR efficiency, the donor template contained a puromycin selection cassette flanked by 500 bp homology arms, one of which housed the G1 variants rs73885319 and rs60910145 (Figure 1A). Successful G1-variant knock-in and cassette removal were confirmed by Sanger sequencing (Figure 1A, Figure S1A-C). Because CRISPR-mediated genome editing can occasionally induce chromosomal changes^22^, we verified that our genome-edited iPSCs maintained a normal karyotype (Figure S1D-E).

To model the G1 variants within a kidney context, we differentiated the G1 and G0 lines into kidney organoids using an adherent culture protocol we have previously established (Figure 1B)^5–7,13,14^. Both G0 and G1 iPSC lines differentiated into kidney organoids without major structural differences, expressing markers of nephron structure including PODXL in glomerular epithelial cells, LTL in the proximal tubule, and E-cadherin in the distal tubule, in appropriately patterned and contiguous segments (Figure 1C).

We next evaluated whether the organoids expressed *APOL1*. RT-qPCR on RNA from whole-well organoids differentiated from G0 and G1 1016SevA lines and the G0 Penn134-61-26 line revealed that organoids expressed little to no *APOL1* under standard culture conditions (Figure 2A-B, Figure S2). IFN-γ has been implicated as a factor that may induce APOL1 expression in mice, although whether this occurs in humans remains unclear^23^. Robust *APOL1* expression (mean >4,000-fold over *ACTB, P* = 0.004 by 2-way ANOVA) was induced in both G0 and G1 organoids when exposed to 25 ng/mL IFN-γ for 24 hours (Figure 2A). APOL1 protein expression was confirmed on immunoblot of whole-well organoids, with the appearance of a ∼40 kDa band, the expected size for APOL1, only in the IFN-γ samples (Figure 2B, Figure S2).

**Figure 2.**
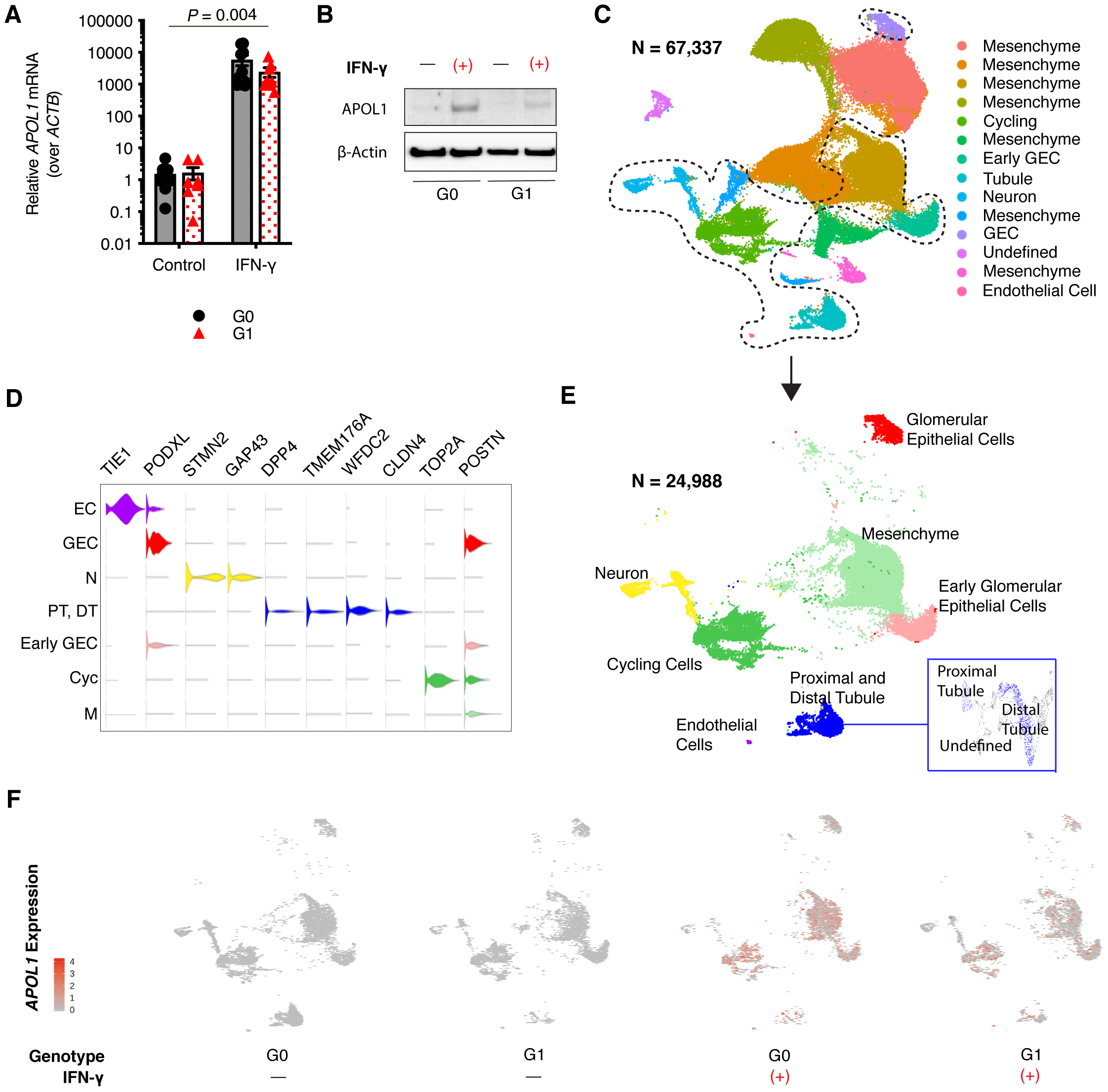
IFN-γ induces APOL1 expression in iPSC-derived kidney organoids. **A.** RT-qPCR of RNA isolated from whole-well G0 and G1 organoids reveal low endogenous APOL1 expression but a greater than 4,000-fold induction of *APOL1* mRNA relative to *ACTB* after 24 hours of 25 ng/mL IFN-γ treatment (*P* = 0.004 by 2-way ANOVA). **B.** Immunoblot of APOL1 expression in G0 and G1 organoids + 25 ng/mL IFN-γ for 24 hours. **C.** UMAP visualization of all whole-well G0 and G1 organoid cells profiled by scCRNA-seq, integrated using Seurat v3. **D.** Violin plot of marker genes used for cluster identification of cell types, color-coded according to labeling of nano-dissected UMAP in **E**, which separates out the main cell types seen in the organoid away from the extra mesenchyme captured in whole-well sequencing. **F.** UMAP feature plots of nano-dissected clusters show *APOL1* expression (red dots) at baseline or when treated with 25 ng/mL IFN-γ for 24 hours. Tun = tunicamycin, EC = endothelial cells, GEC = glomerular epithelial cells, N = neuron, PT = proximal tubule, DT = distal tubule, Cyc = cycling, M = mesenchyme.

To reveal which organoid cell types express *APOL1*, and underlying effects of *APOL1* risk variants on gene expression, we performed scRNA-seq on all cells collected from whole wells of G0 and G1 organoids from three separate experiments. A total of 67,337 cells across genotype and experimental conditions passed quality-control metrics (see Supplemental Methods) and were analyzed using the integrated analysis workflow of Seurat v3^24^ (Figure 2C), which was chosen to be the primary analysis pipeline to improve normalization for batch effects and evaluate differential expression across clusters. To further confirm our findings, we also analyzed the data separately with an unsupervised deep embedding algorithm for single-cell clustering (DESC)^25^. Primary output of unsupervised clustering by Seurat v3 yielded 13 clusters (Figure 2C, S3A), seven of which were determined to consist of mesenchymal cells, while the others included glomerular epithelial cells at early and more mature stages, proximal and distal tubule, cycling cells, neurons, and endothelial cells (Figure 2D-E, Figure S3A) as determined by canonical marker genes (Figure 2D). The abundance of mesenchyme versus epithelial lineages in these whole-well differentiations is high but within the expected range for different iPSC lines and batches of organoids^7,14^. These clusters were present in both G0 and G1 organoids (Figure S3B). Similar unsupervised clustering was also obtained using DESC (Figure S4A-B). Like the RT-qPCR and immunoblot assays, scRNA-seq revealed that resting organoids do not express *APOL1*. However, exposure to IFN-γ for 24 hours induced *APOL1* expression across all cell types, including glomerular epithelial cells, endothelial cells, and tubular cells (Figure 2F, Figure S5).

In contrast to an inducible *APOL1* risk-variant overexpression cell model^19^, G1 kidney organoids did not undergo appreciable cell death when *APOL1* was expressed (Figure S3B). To determine whether risk-variant *APOL1* expression causes transcriptome-wide changes, we performed differential expression analysis using Seurat v3. Comparing G0 and G1 organoids exposed to IFN-γ, we found greater variance in gene expression patterns when examining each cell type separately than when examining the whole organoid (Figure 3A). In IFN-γ stimulated organoids, very few genes differentially expressed between G0 and G1 glomerular epithelial cells overlapped with genes differentially expressed between G0 and G1 tubular cells (Figure 3B), suggesting that risk-variant *APOL1* could potentially alter transcriptional programs in a cell-type specific fashion. This result is concordant with findings that human genetic variation exerts cell-type specific effects in CKD^11^ and with a prior finding that glomerular and tubular compartments of APOL1-mediated focal segmental glomerulosclerosis biopsies exhibit distinct gene expression patterns^26^.

**Figure 3.**
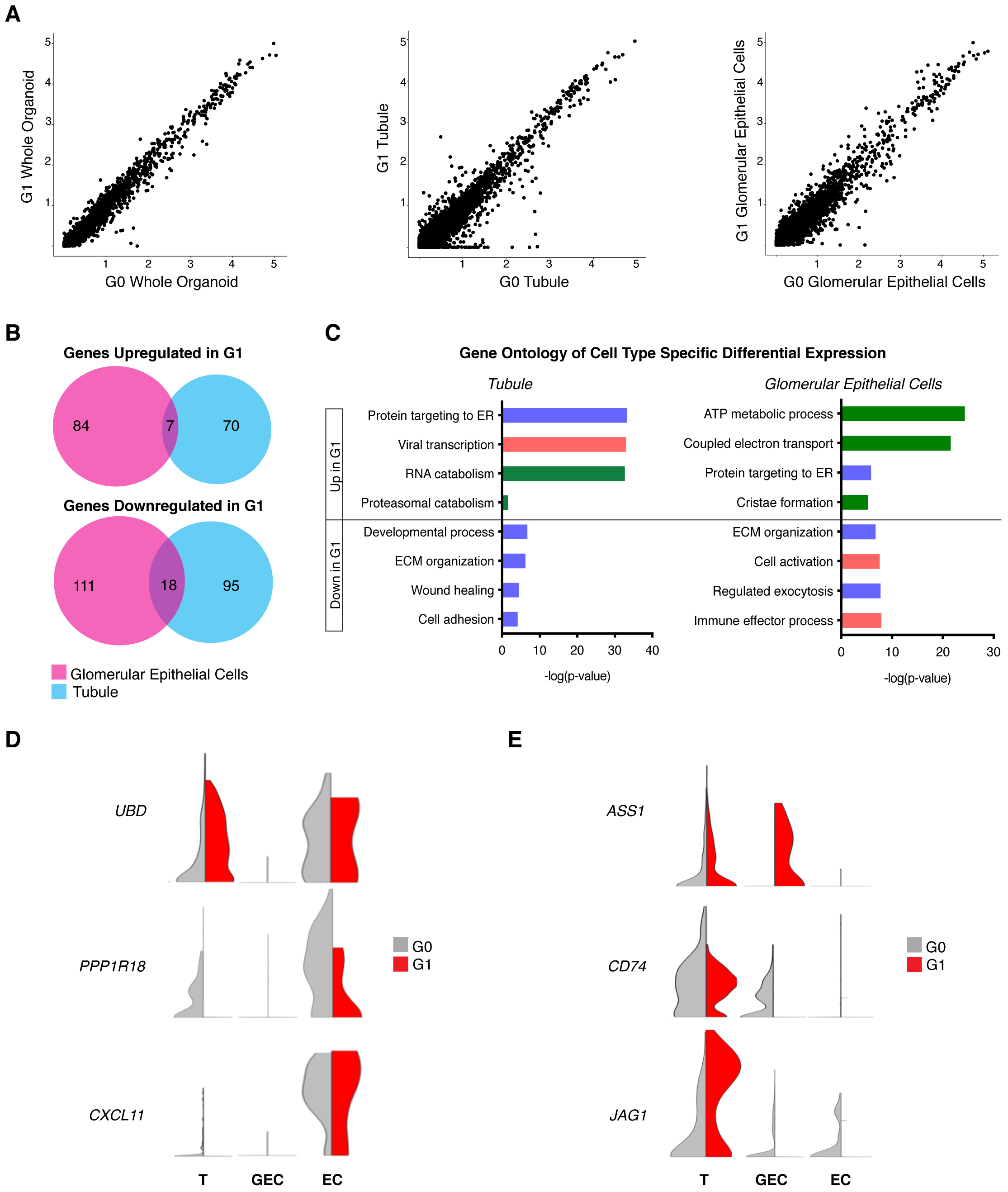
scRNA-seq reveals cell-type specific differential expression patterns between G0 and G1 organoids treated with IFN-γ. **A.** Scatter plots of G0 vs. G1 organoid gene expression values across all cell types (left), among tubular cells (middle), and among glomerular epithelial cells (right). Greater variance is seen in the cell-type specific plots. **B**. Venn diagrams show little overlap of expression patterns across cell types when comparing which genes are upregulated in G1 IFN-γ stimulation over G0 IFN-γ stimulation. Similarly low overlap across cell types for genes downregulated in G1 IFN-γ stimulation. **C.** Gene ontology of the differentially expressed genes within each cell type, with -log(p-value) plotted for each biological process. Metabolic (green), stress (pink), cell and organelle function (purple), collagen / matrix (grey). **D.** Violin plots visualizing cell type-specific expression patterns of putative regulators ^18,26,27^ of APOL1 abundance. **E.** Violin plots visualizing cell type-specific expression patterns of novel genes.

Indeed, the genes differentially expressed between G0 and G1 glomerular epithelial cells stimulated with IFN-γ were enriched for biological processes related to metabolic function, whereas tubular cells stimulated with IFN-γ exhibited differential (between G0 and G1) expression of genes related to inflammation and protein targeting and processing (Figure 3C). To visualize these cell-type specific gene expression patterns induced by IFN-γ, we generated normalized transcript abundance on violin plots for each cluster (Figure 3D-E). We first plotted genes previously identified by other studies to be potential modifiers of APOL1-mediated kidney disease. A large linkage disequilibrium block on chromosome 6 containing *UBD* and *PPP1R18* may house disease-modulating genes ^26,27^, with visible differences between *APOL1* genotypes in *UBD* and *PPP1R18* expression in the tubule and endothelial clusters (Figure 3D). *CXCL11* was previously found to be upregulated in the glomeruli of patients with APOL1-mediated kidney disease ^26^; it appears mildly more abundant in G1 endothelial cells (Figure 3D). Novel genes such as *ASS1* are specifically upregulated in interferon-stimulated G1 glomerular epithelial cells, while *CD74* and *JAG1* are markedly downregulated in G1 glomerular epithelial cells compared to G0 cells (Figure 3E).

With these cell-type specific differences in G1 organoid gene expression, we next evaluated whether introducing an additional stressor alters G1 cell phenotypes. Because APOL1-mediated kidney disease is incompletely penetrant, others have proposed a “second-hit hypothesis” that individuals carrying two *APOL1* risk alleles develop disease after exposure to environmental or genetic modifiers ^28,29^. We tested whether ER stress could provide a stimulus to alter stress response and cellular phenotypes in the high-risk genotype. ER stress was chosen because ubiquitin D, encoded by *UBD*, is involved in the ER stress response, and risk-variant APOL1 appears to localize to the ER membrane^30^. Furthermore, ER stress is becoming an increasingly recognized driver of complex diseases, including CKD ^31–34^. To induce ER stress, we treated organoids with 5 µM tunicamycin for 24 hours and observed increased expression of unfolded protein response genes *EDEM1* and *HSPA5* by RT-qPCR (Figure 4A). Violin plots of the podocyte marker *PODXL* in organoids treated with IFN-γ and tunicamycin revealed relatively decreased expression of *PODXL* in the G1 glomerular epithelial cell cluster, suggesting the G1 organoid may acquire a less differentiated state under stress (Figure 4B). We also discovered that G1 organoids treated with both IFN-γ and tunicamycin demonstrated less distinct cluster topology seen in partition-based graph abstraction (PAGA)^35^. More specifically, G1 glomerular epithelial cells became closer to the other cell types in spatial relationship, whereas G0 glomerular epithelial cells still remained more distinct from the other cell clusters (Figure 4C). Trajectory inference of G0 and G1 organoid glomerular epithelial cells (Figure 4D) revealed that G1 organoids, compared to G0 organoids, subjected to ER stress have more cells scattered along trajectory paths between clusters, as well as a smaller proportion of mature glomerular epithelial cells compared to early glomerular epithelial cells (Figure 4E-F), consistent with potential dedifferentiation of G1 glomerular epithelial cells during stress.

**Figure 4.**
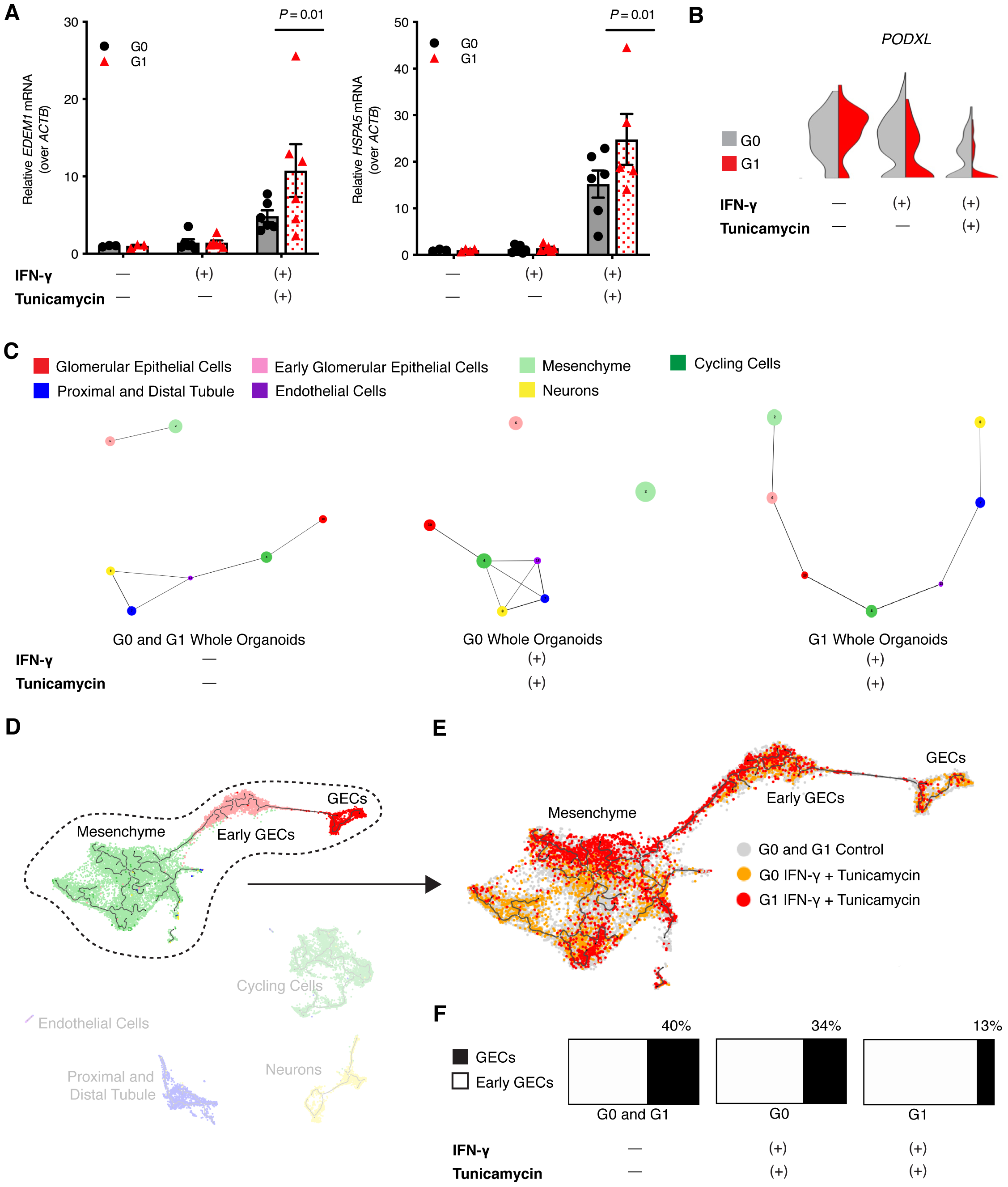
ER stress increases stress response and dedifferentiation of glomerular epithelial cells (GECs) in G1 kidney organoids. **A.** RT-qPCR of RNA isolated from whole-well G0 and G1 organoids reveals increased unfolded protein response gene expression (*EDEM1, HSPA5*) in G1 organoids exposed to both IFN-γ and tunicamycin (*P* = 0.01 by 2-way ANOVA). **B.** Violin plots of *PODXL* expression in the GEC clusters of G0 and G1 organoids subjected to either IFN-γ alone or both IFN-γ and tunicamycin. **C.** Partition-based graph abstraction (PAGA)^35^ visualizes trajectory inference of all organoid single cells. The more mature GECs are connected to non-mesenchyme cell clusters in the topology of the control organoids. For G0 organoids subjected to both IFN-γ and tunicamycin, the more mature glomerular cells are still more distant from the mesenchyme. PAGA visualization of G1 organoids subjected to both IFN-γ and tunicamycin reveals less distinct separation of GECs from early GECs and mesenchyme. **D.** Trajectory inference UMAP of all sequenced organoids was created using Monocle 3. The trajectory of the relationship among mesenchyme, early GECs, and more mature GECs is circled and taken forward to **E**, where G1 organoids treated with both IFN-γ and tunicamycin (red) have a relatively smaller proportion of GECs to early GECs compared to G0 organoids subjected to the same stressors (orange). **F.** The relative number of GECs (black) to early GECs (white) is represented by the bar charts of G0 and G1 control organoids (left), G0 organoids treated with IFN-γ and tunicamycin (middle), and G1 organoids treated with IFN-γ and tunicamycin (right).

In summary, we have conducted profiling of genome-edited iPSC-derived kidney organoids to detect the effect of *APOL1* risk variants. Our approach circumvents the challenges of model organisms by expressing the G1 variants in their native genomic context. In addition, footprint-free CRISPR-Cas9 genome editing allows for interrogation of the risk variants on isogenic backgrounds, reducing genetic and epigenetic heterogeneity as potential confounding modifiers of *APOL1* expression and function. Using single-cell transcriptomics revealed cell-type specific gene expression differences between G0 and G1 organoids when *APOL1* is induced. Furthermore, we demonstrated that additional stressors can lead to more pronounced dedifferentiation of glomerular epithelial cells in G1 organoids, although we recognize the glomerular epithelial cells from these iPSC lines are relatively immature. Our work will require further validation from additional risk-variant *APOL1* iPSC lines with matching isogenic controls. However, our model system, which combines the power of footprint-free genome editing with organoid culture, provides a human-relevant platform to launch larger-scale mechanistic and screening studies for APOL1-mediated kidney disease.

## AUTHOR CONTRIBUTIONS

Manuscript concept developed by J.L. and B.S.F. Data collected by E.L., D.Y., A.D., and B.H.C. Bioinformatics analysis was performed by B.R. and M.L. Manuscript was drafted by J.L., E.L., B.R., and B.S.F. and edited by A.D., M.D.D., K.D.S., and J.H. Final edits were made by J.L. and B.S.F. All authors approved the final version of the manuscript.

## ACKNOWLEDGEMENTS

The views expressed in this article are those of the authors and do not necessarily reflect the position or policy of the Department of Veterans Affairs or the United States government. We thank Dr. Kiran Musunuru for his input on footprint-free CRISPR Cas-9 genome editing. We appreciate the massively parallel sequencing services provided by the University of Illinois Genomics Core. A portion of this work was supported by the National Institutes of Health grants K08 HL135348 (to J.L.) and UG3TR002158 (to J.H.).

