## Supplement for "Profiling *APOL1* Nephropathy Risk Variants in Genome-Edited Kidney Organoids with Single-Cell Transcriptomics"

**SUPPLEMENTAL TABLE OF CONTENTS**

**Figure S1.** CRISPR-Cas9 engineering of G0 and G1 induced pluripotent stem cells (iPSCs) on isogenic backgrounds.

**Figure S2.** APOL1 protein expression induced by IFN-$\gamma$.

**Figure S3.** UMAP visualization by cell-type cluster and by experimental conditions.

**Figure S4.** Scanpy visualization of organoid scRNA-seq analyzed with the DESC pipeline.

**Figure S5.** *APOL1* mRNA expression induced by IFN-$\gamma$.

**Figure S6**. Original uncropped images of Western blots for APOL1 protein expression in whole-well kidney organoid samples.

**Table S1.** Oligonucleotides

**Supplemental Methods: Details of single-cell RNA-seq (scRNA-seq) analyses**

**Supplemental References: Specific to text in this supplement**

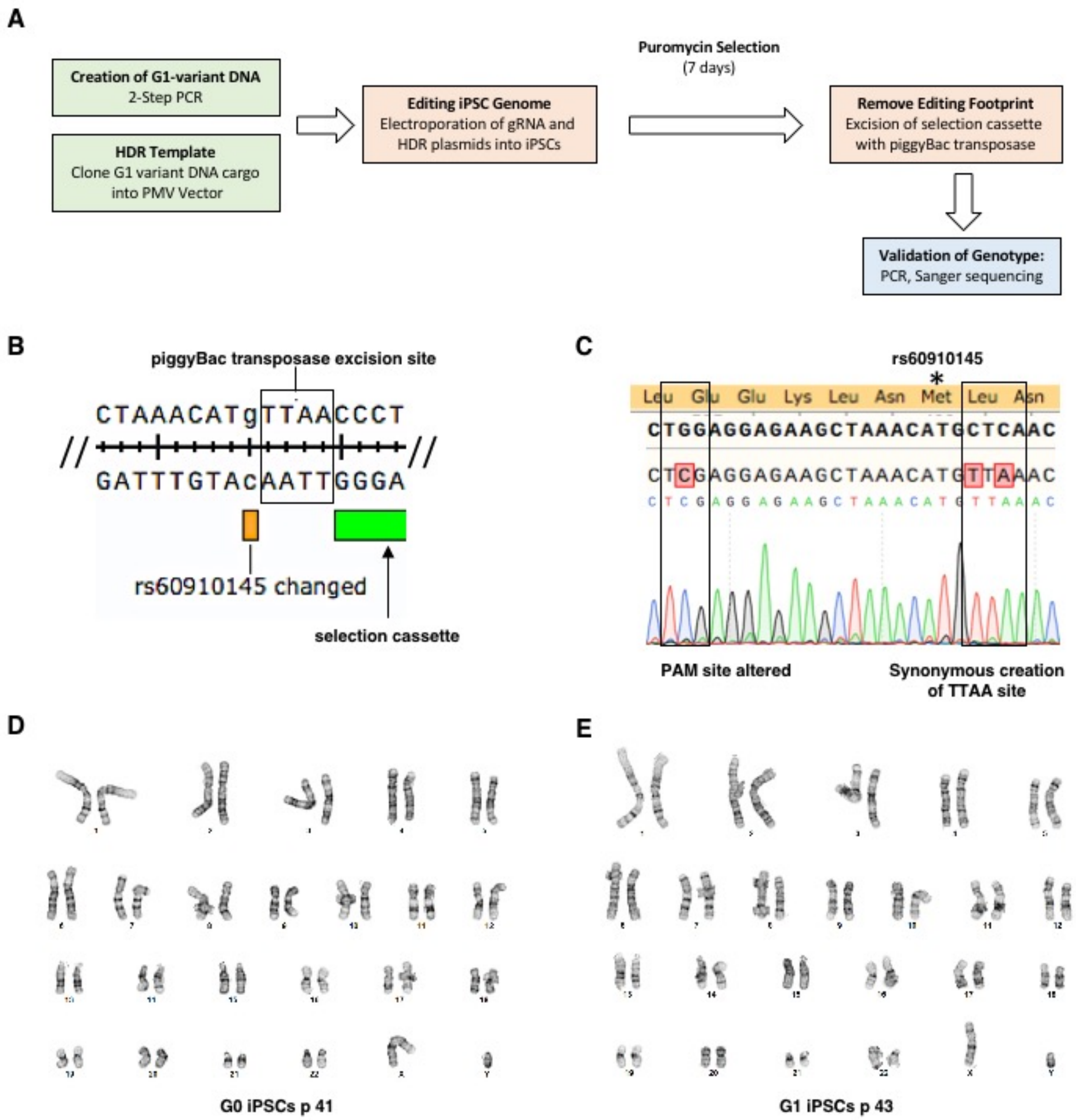


**Figure S1.** CRISPR-Cas9 engineering of G0 and G1 induced pluripotent stem cells (iPSCs) on isogenic backgrounds. **A.** Overview of the workflow for knocking in the G1 variants into the 1016SevA iPSC line. **B.** Partial sequence of homology-directed repair (HDR) donor template at junction of one G1 SNP (rs60910145) flanking the synonymous introduction of a TTAA piggyBac transposase excision site adjacent to the puromycin selection cassette. **C.** Sanger sequencing validation of synonymous sequence changes introduced to facilitate “scarless” knock-in of the G1 variants. The left black rectangle highlights a synonymous G to C change that alters the PAM site to prevent re-cutting of a successfully knocked in template. The right black rectangle highlights the synonymous change of a CTC (Leu) to TTA (Leu) to facilitate the introduction of the TTAA piggyBac excision site. **D.** Karyotype of the 1016SevA G0 iPSC line. **E.** Karyotype of the 1016SevA G1 iPSC line.

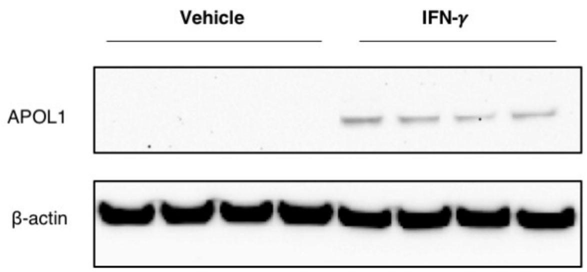


**Figure S2**. *APOL1* protein expression induced by IFN-γ. Immunoblot for APOL1 protein expression in whole-well organoids differentiated from the Penn134-61-26 iPSC line (G0 genotype, African ancestry).

**A**

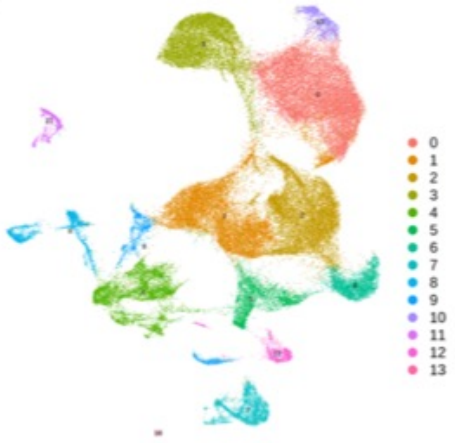


| Cell Type | Cluster ID | % of Total Cells |
|---|---|---|
| GEC | 10 | 1.8 |
| Early GEC | 6 | 4.2 |
| Tubule | 7 | 3.3 |
| Endothelial Cell | 13 | 0.2 |
| Neuron | 8 | 2.9 |
| Cycling | 4 | 7.8 |
| Mesenchyme | 0-3, 5, 9, 12 | 78.4 |
| Artifact | 11 | 1.4 |

**B**

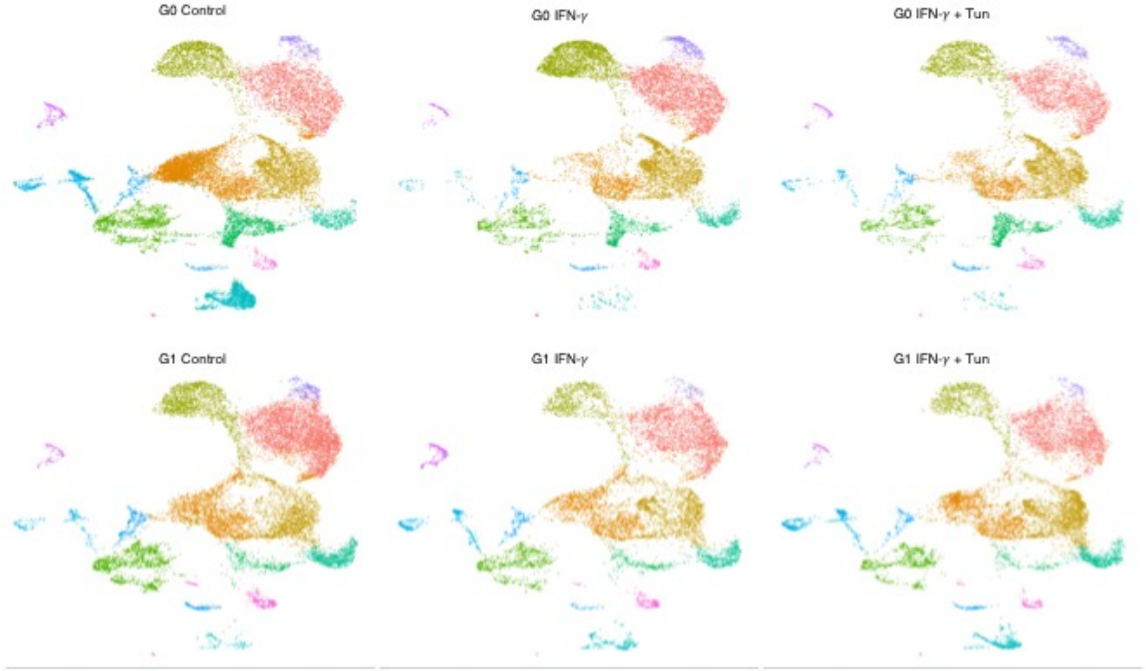


**Figure S3.** UMAP visualization by cell-type cluster and by experimental conditions. **A.** UMAP visualization of integrated dataset of sequenced organoids, analyzed with Seurat v3. 13 cell-type clusters were identified. Percentages of each cell type among all sequenced cells are listed in the table on the right. **B.** UMAP plots of organoids from each separate genotype and experimental condition show representation of each cell type in each condition.

**A**

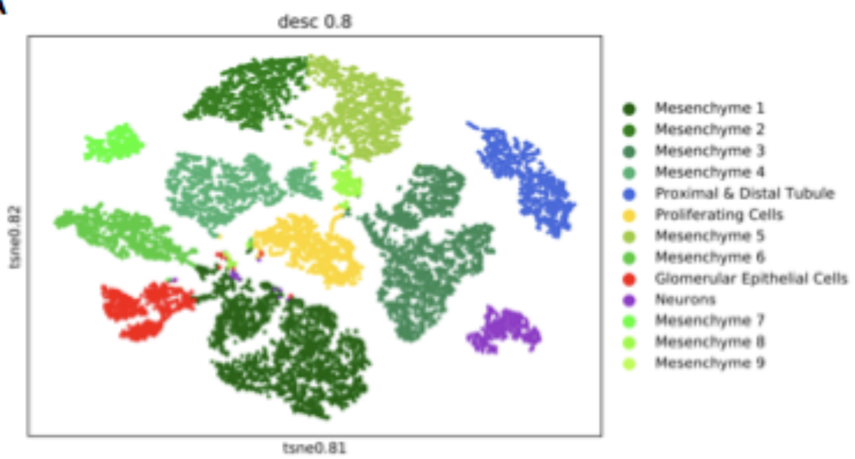


**B**

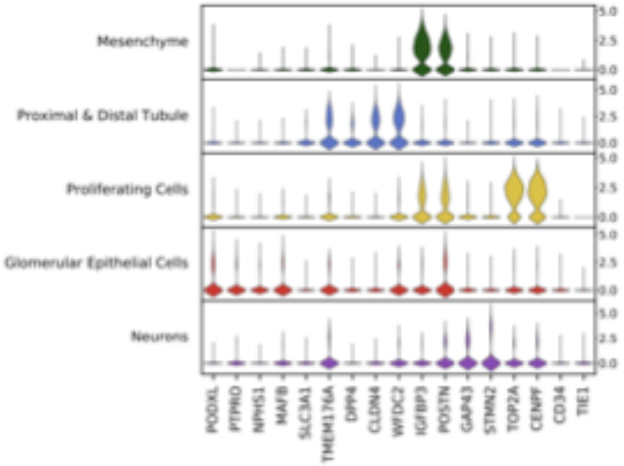


**Figure S4.** Scanpy visualization of organoid scRNA-seq analyzed with the DESC pipeline. **A.** All sequenced organoids represented on t-SNE plot with 13 cell-type clusters identified. **B.** Violin plot of marker genes of identified cell-type clusters.

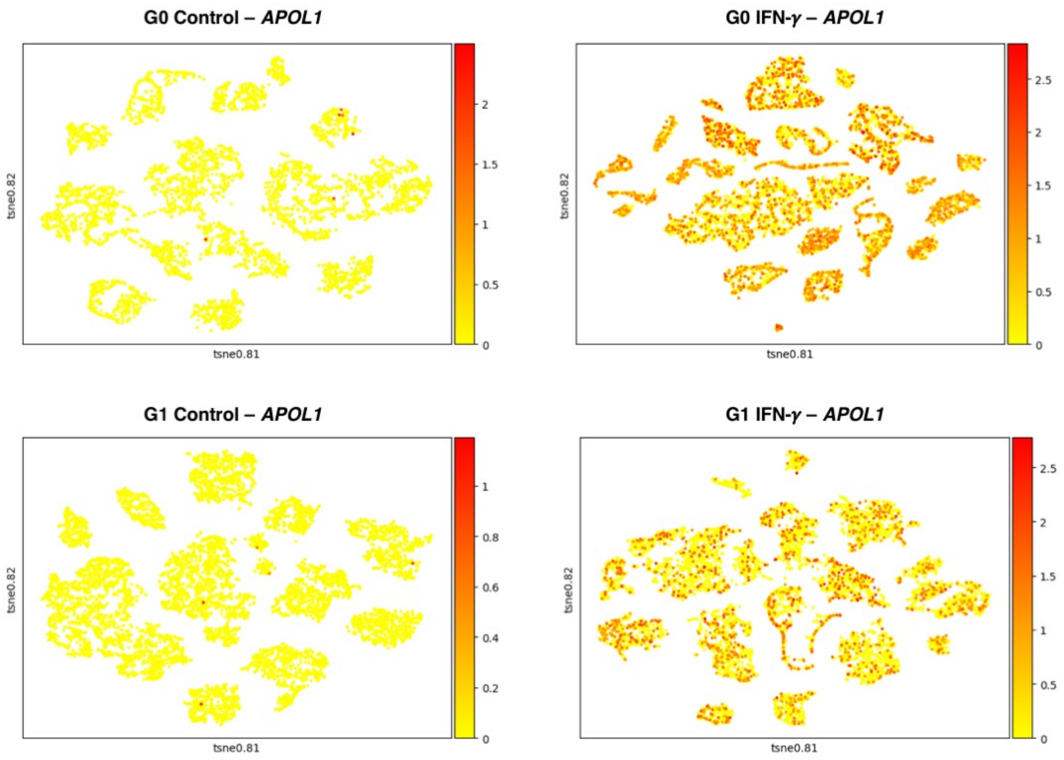


**Figure S5.** *APOL1* mRNA expression induced by IFN-γ. Scanpy feature plots of *APOL1* expression in whole-well organoids as analyzed by the DESC pipeline. Cells with *APOL1* expression are visualized in red.

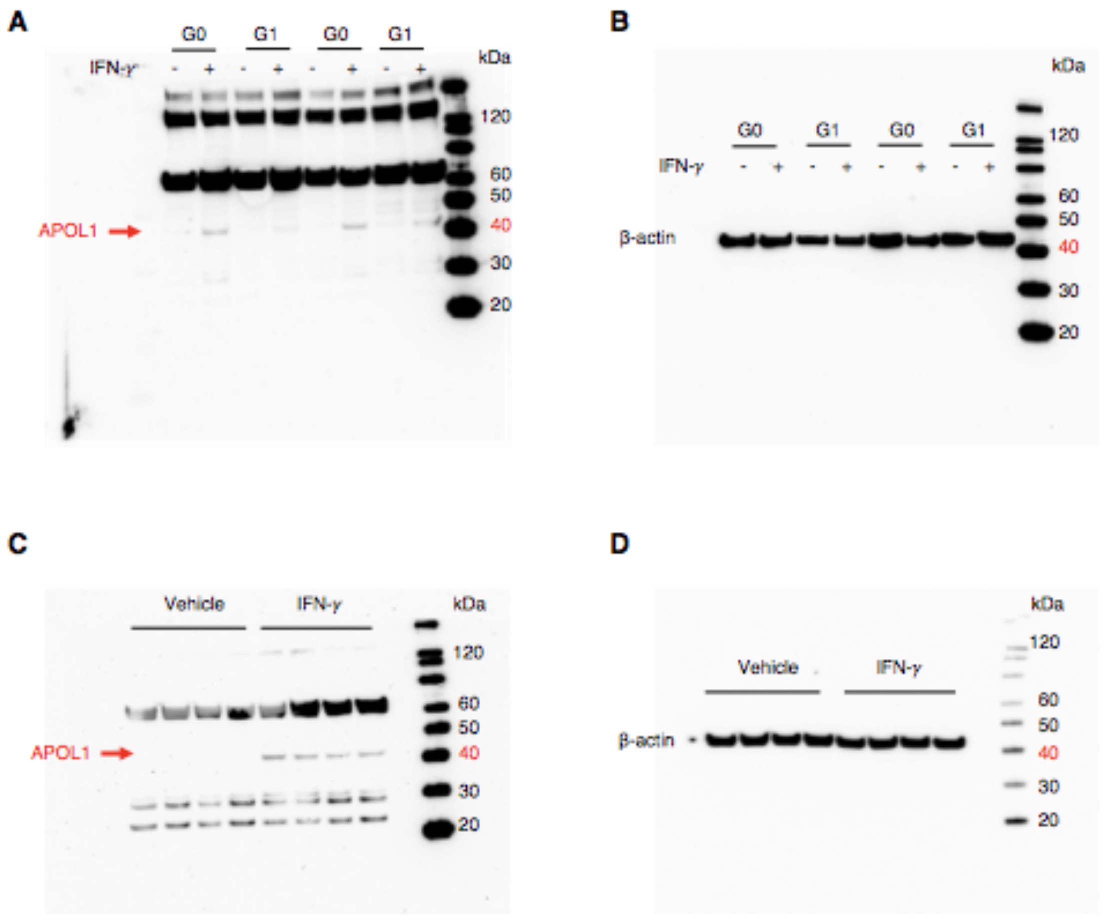


**Figure S6.** Original uncropped images of Western blots for APOL1 protein expression in whole-well kidney organoid samples. **A.** APOL1 expression in whole wells of kidney organoids differentiated from the 1016SevA G0 and G1 lines subjected to 24 hours of vehicle or 25 ng/mL IFN-γ. APOL1-specific bands (red arrow) at the expected 44 kDa size. **B.** Immunoblot of beta-actin of the same samples seen in (**A**). **C.** APOL1 expression in whole wells of kidney organoids differentiated from the Penn134-61-26 iPSC line (G0 genotype) subjected to 24 hours of vehicle or 25 ng/mL IFN-γ. APOL1-specific bands (red arrow) at the expected 44 kDa size. **D.** Immunoblot of beta-actin of the same samples seen in (**C**).

**Table S1. Oligonucleotides**

| Oligonucleotide Description | 5' to 3' Sequence |
|---|---|
| Amplification of *APOL1* gDNA to create template of homology arm A upstream of selection cassette - forward primer | GGCAGCCTTGTACTCTTGGA |
| Amplification of *APOL1* gDNA to create template of homology arm A upstream of selection cassette - reverse primer | GCCCTGTGGTCACAGTTCTT |
| Amplification of *APOL1* gDNA to create template of homology arm B downstream of selection cassette - forward primer | GAGCTGGAGGAGAAGCTAAACA |
| Amplification of *APOL1* gDNA to create template of homology arm B downstream of selection cassette - reverse primer | TTCTCCTTGCTGCACTGCCA |
| Homology arm A insert for TA cloning - forward primer | TGCGGCCGCCCGCTTTGACCGGGATTACC |
| Homology arm A insert for TA cloning - reverse primer | TGGTCTCTTTAAAATGTTTAGCTTCTCCTCGAGCTCCTGAGCCACCTTC |
| 2-step PCR introduction of G1 variants - forward primer 1 | TGCGGCCGCCCGCTTTGACCGGGATTACC |
| 2-step PCR introduction of G1 variants - reverse primer 1 | CCAGCACAAGAAAGAAGCcTACAGGGGCCACATCCGTG |
| 2-step PCR introduction of G1 variants - forward primer 2 | CACGGATGTGGCCCCTGTAgGCTTCTTTCTTGTGCTGG |
| 2-step PCR introduction of G1 variants - reverse primer 2 | TGGTCTCTTTAAcATGTTTAGCTTCTCCTCgAGCTCCTGAGCCACCTTC |
| Homology arm B insert for TA cloning - forward primer | TGAAGACAATTAAACAATAATTATAAGATTCTG |

| | |
|---|---|
| Homology arm B insert for TA cloning<br>- reverse primer | TCCATGGCCTCCCCTGCTGTGCTCAGC |
| Guide RNA oligo #1 | CACCGGAAGAAGGTGGCTCAGGAGC |
| Guide RNA oligo #2 | AAACGCTCCTGAGCCACCTTCTTCC |

### SUPPLEMENTAL METHODS

#### Processing of 10x Genomics Chromium scRNA-seq data

Raw scRNA-seq data were processed using the 10x Genomics CellRanger software (version 1.3.0). BCL files obtained from the Illumina NovaSeq platform were converted into Fastq files using the Cell Ranger *mkfastq* program. Fastq files were then mapped to the hg38 reference, downloaded from the 10x Genomics website (http://cf.10xgenomics.com/supp/genome/refdata-GRCh38-2.1.0.tar.gz). Genes were counted using the CellRanger *count* program. Because 5' chemistry libraries were generated, the "SC5P-PE" option was used for the chemistry parameter to capture full-length paired end reads for more depth. The *aggr* program was run to generate aggregate count matrices for replicate libraries of the same genotype and experimental condition.

#### Filtering, dimensionality reduction, and clustering of scRNA-seq data (primary workflow)

Processing of scRNA-seq data was performed in R[1] using the Seurat v3 package[2]. Initially each of the individual samples were converted into a Seurat object after removing any genes expressed in fewer than five cells and any cells expressing fewer than 300 detected genes. Cells with over 25% raw unique molecular identifiers (UMIs) mapping to mitochondrial genes were filtered out, as were control cells with over 4,000 RNA feature counts, and stimulated cells with over 6,000 RNA feature counts based on quality control graphs. These objects were normalized using the default *LogNormalize* method and their variable features identified using variance-stabilizing transform. The integrated analysis pipeline was then used to combine the count matrices from all libraries into one single Seurat object by first identifying the integration anchors and then providing those anchors to the *IntegrateData* function. The integrated object was scaled using the *ScaleData* function and its dimensions with the most variance identified using *RunPCA*. Lastly, a combined UMAP plot was created (*RunUMAP*) and the cell clusters identified (*FindClusters*) by using the first twenty dimensions and allowing for a resolution of 0.2.

#### Filtering, dimensionality reduction, and clustering of scRNA-seq data (secondary workflow)

DESC[3] employs many of the same steps used by Seurat v3, with the exception of using Tensorflow machine learning to cluster the various cell types. Count files from CellRanger were filtered by removing cells with fewer than 200 genes and genes expressed in fewer than 3 cells. Cells with greater with greater than 10% raw UMIs mapping to mitochondrial genes and with greater than 3,500 total RNA features were removed. Cells were normalized using the *LogNormalize* function and log transformed using the *log1p* function. Highly variable genes were identified using the values, 0.0125, 3 and 0.5 for the parameters "min_mean", "max_mean" and "min_disp", respectively. Finally, the data were scaled with the *scale* function and the clusters were identified using the *desc.train* function.

#### Trajectory Inference

The latest version of Scanpy's PAGA[4] and Monocle 3[5] were used to identify the trajectories from the various cell types. Clusters that had been identified using Seurat v3 were preserved and applied into both Monocle and PAGA. Thus, the steps involving the classification and clustering of the cells was performed but overwritten using Seurat v3 clusters. The Monocle 3 cell_data_set (CDS) object was created by extracting the single-cell gene expression, cell metadata, and gene identification information from a Seurat object, combining them into a CDS object with the new_cell_data_set function. Prepossessing was conducted on the CDS object by normalizing the data using the first 100 principal components. Each UMI was plotted on a UMAP plot by having dimensions reduced using the 'reduce_dimension' function. These UMIs were then grouped into Louvain clusters and PAGA partitions (cluster_cells) followed by the creation of a principal graph to visualize the trajectory for each of the cells (learn_graph). The same cell clusters from Seurat v3 were inserted into PAGA to perform topology analysis, which identifies connected, disconnected, and cyclical graphs in addition to the ones generated by Monocle. The analysis was started by identifying the top principal components for the cell clusters identified by Seurat using the 'arpack' method. A nearest neighbor analysis was then performed with the 'n_neighbors' parameter set to 15 and using the first 40 principal components. Finally, a PAGA plot was created using the Seurat clusters and a threshold value of 0.5 to visualize the more significant connections between cell cluster nodes.

**Gene Ontology**

Gene ontology for differentially expressed genes was performed using the PANTHER[6] classification system's statistical overrepresentation test against all coding human genes.
